# Sex-specific gene and pathway modeling of inherited glioma risk

**DOI:** 10.1101/235408

**Authors:** Quinn T. Ostrom, Warren Coleman, William Huang, Joshua B. Rubin, Justin D. Lathia, Michael E. Berens, Gil Speyer, Peter Liao, Margaret R. Wrensch, Jeanette E Eckel-Passow, Georgina Armstrong, Terri Rice, John K. Wiencke, Lucie S. McCoy, Helen M. Hansen, Christopher I. Amos, Jonine L. Bernstein, Elizabeth B. Claus, Dora Il’yasova, Christoffer Johansen, Daniel H. Lachance, Rose K. Lai, Ryan T. Merrell, Sara H. Olson, Siegel Sadetzki, Joellen M. Schildkraut, Sanjay Shete, Richard S. Houlston, Robert B. Jenkins, Ulrika Andersson, Preetha Rajaraman, Stephen J. Chanock, Martha S. Linet, Zhaoming Wang, Meredith Yeager on behalf of the GliomaScan consortium^, Beatrice Melin, Melissa L. Bondy, Jill S. Barnholtz-Sloan

## Abstract

**Background:** Genome-wide association studies (GWAS) have identified 25 risk variants for glioma, which explain ~30% of heritable risk. Most glioma histologies occur with significantly higher incidence in males. A sex-stratified analysis ide7ntified sex-specific glioma risk variants, and further analyses using gene- and pathway-based approaches may further elucidate risk variation by sex.

**Methods:** Results from the Glioma International Case-Control Study were used as a testing set, and results from three GWAS were combined via meta-analysis and used as a validation set. Using summary statistics for autosomal SNPs found to be nominally significant (p<0.01) in a previous meta-analysis and X chromosome SNPs with nominally significant association (p<0.01), three algorithms (Pascal, BimBam, and GATES) were used to generate gene-scores, and Pascal was used to generate pathway scores. Results were considered significant when p<3.3x10^−6^ in ⅔ algorithms.

**Results:** 25 genes within five regions and 19 genes within six regions reached the set significance threshold in at least 2/3 algorithms in males and females, respectively. *EGFR* and *RTEL1-TNFRSF6B* were significantly associated with all glioma and glioblastoma in males only, and a female-specific association in *TERT*, all of which remained nominally significant after conditioning on known risk loci. There were nominal associations with the Telomeres, Telomerase, Cellular Aging, and Immortality pathway in both males and females.

**Conclusions:** These results suggest that there may be biologically relevant significant differences by sex in genetic risk for glioma. Additional gene- and pathway-based analyses may further elucidate the biological processes through which this risk is conferred.

## INTRODUCTION

Glioma is the most common type of primary malignant brain tumor in the United States (US), with an average annual age-adjusted incidence rate of 6.0/100,000.^1^ Glioma can be broadly classified into glioblastoma (GBM, 61.9% of gliomas in adults 18+ in the US) and lower-grade glioma (non-GBM glioma, 24.2% of adult gliomas).^1^. Gliomas are significantly more common in people of European ancestry, in males and in older adults.^1^. Most glioma histologies occur with a 30-50% higher incidence in males, and this male preponderance of glial tumors increases with age in adult glioma (**Supplemental Figure 1**).^1^

Many environmental exposures have been investigated as sources of glioma risk, but the only validated risk factors for these tumors are ionizing radiation (which increases risk), and history of allergies or other atopic disease (which decreases risk).^2^ The contribution of common low-penetrance SNPs to the heritability of sporadic glioma in persons with no documented family history is estimated to be ~25%.^3^ A recent glioma genome-wide association study (GWAS) meta-analysis validated 12 previously reported risk loci, and identified 13 new risk loci, and these 25 loci in total are estimated to account for ~30% of heritable glioma risk.^4^ This suggests that there are both undiscovered environmental risk (which accounts for ~75% of incidence variance) and genetic risk factors (accounting for ~70% of heritable risk).^3,4^ Each individual GWAS results in regression estimates for hundreds of thousands of single nucleotide polymorphisms (SNPs), only several hundred of which may be prioritized for further investigation. While this process is appropriate for identifying individual loci that contribute to the development of disease, there is likely additional information about disease risk within results that do not meet thresholds for statistical significance. Single-SNP tests may not be appropriate for additional loci discovery given the known biological complexity of gliomas. Multi-SNP methods, such as gene or pathway-based approaches, can allow for additional discovery in a manner that complements single-SNP approaches, while substantially reducing the multiple testing burden associated with GWAS.^5^ As an example, an association study focused on the cAMP pathway identified SNPs in Adenylate Cylase 8 as a sex-specific modifier of risk for low grade astrocytoma in Neurofibromatosis Type 1.^6^ A more recent sex-stratified analysis also identified glioma risk loci that differ by sex.^7^ Additional analyses using gene- and pathway-based approaches may further elucidate sex differences in genetic risk for glioma.

## METHODS

Using summary statistics for autosomal markers found to be nominally significant (p<0.01) in a previous 8-study meta-analysis^8^ (**Figure 1a**) and X chromosome makers with nominally significant single SNP association (p<0.01), three algorithms, Pascal,^9^ BimBam,^10^ and GATES,^11^ were used to generate gene-scores. Gene-based effects were assessed using all SNPs within 50kb of each gene (using 5’ and 3’ UTR) as defined using the UCSC hg19 assembly. Results from the Glioma International Case-Control Study (GICC)^8,12^were used as a testing set (**Figure 1a**), and results from three prior glioma GWAS (San Francisco Adult Glioma Study GWAS,^13^ MD Anderson Glioma GWAS,^14^ and National Cancer Institute’s Gliomascan^15^) were combined via inverse-variance weighted fixed effects meta-analysis in META^16^ and used as a validation set for any significant genes and pathways (**Figure 1a**). See **Supplemental Table 1** for an overview of characteristics for individuals included in these datasets, and **Figure 1** for an overview of the study schematic.

**Figure 1.**
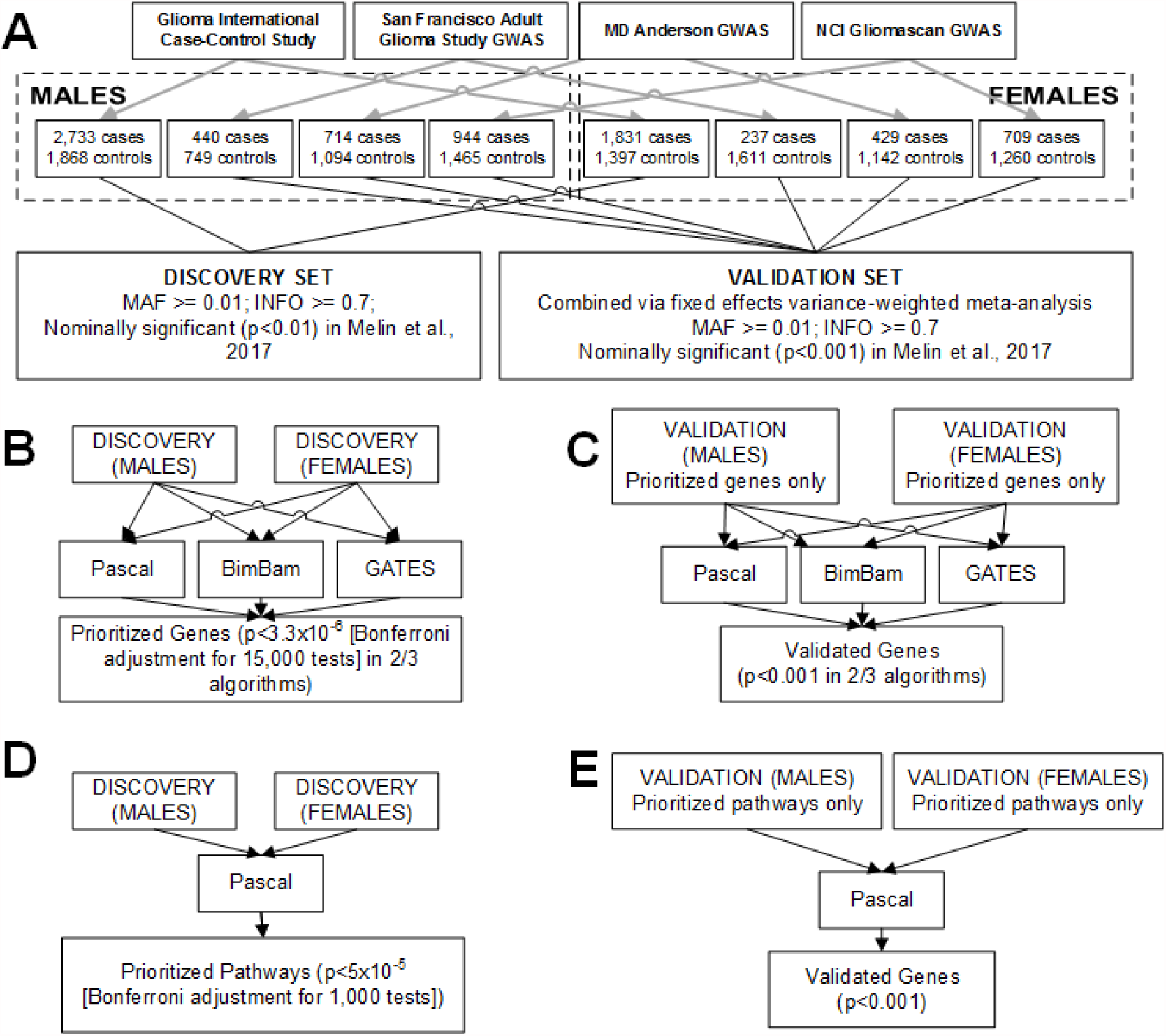
*Study schematic for a) generation of discovery and validation summary statistic sets, b) generation of discovery gene-based tests and prioritization c) validation of gene-based tests, d) generation of discovery pathway-based tests and prioritization e) validation of pathway-based tests*.

Summary statistics were generated using sex-stratified logistic regression models in SNPTEST.^17^ Autosomal chromosomes were analyzed using sex-stratified logistic regression models to estimate sex-specific betas (β_M_ and β_F_), standard errors (SE_M_ and SE_F_), and p values (p_M_ and p_F_). X chromosome data were available from GICC set only, and analyzed using logistic regression model in SNPTEST module ‘newml’ assuming complete inactivation of one allele in females, and males are treated as homozygous females. Linkage disequilibrium (LD) information was based on structure within the European cases from the 1,000 genomes project phase 3 dataset. All analyses were performed separately for males and females to identify genes and pathways with germline variation between cases and controls. Genes were prioritized that were identified by at least 2 of the 3 selected algorithms (**Figure 1b-c**). Analyses were conducted for glioma overall, and for glioblastoma only, by sex within each dataset.

Pascal^9^ calculates gene scores using the VEGAS^18^ scoring algorithm, generates a gene-based test statistic using sum-of-chi-squares (SOCS) correcting for linkage disequilibrium (LD) structure (based on a reference set). Genes that are in LD are considered to be a ‘fusion gene’ and have only one gene score calculated. Bimbam^10^ (as implemented in FAST using summary statistics^19^) is a Bayesian regression approach. This method calculates an average Bayes Factor for all K possible models within a gene, where K is the number of SNPs. The model then uses a Laplace method to estimate posterior distributions of the model’s parameters, and distribution models are obtained using the Fletcher-Reeves conjugate gradient algorithm. GATES^11^ (as implemented in FAST^19^) uses a modified Sims test that combines SNP-based p values, using the p value correlation matrix to estimate the number of independent SNPs within the gene. The resulting gene-based p values approximate a uniform distribution. For all methods implemented within FAST, SNPs were excluded if they were in complete LD (r^2^=1) with another SNP in the gene, which limited the amount of SNPs evaluated within each gene.

Pathway scores were generated using Pascal,^9^ using gene and fusion-gene scores generated by the Pascal algorithm (**Figure 1d-e**). The pathway score was then calculated using both independent and fusion genes. A parameter free enrichment strategy was used to calculate pathway scores using either a chi-squared method (gene score p values were ranked and transformed to a uniform distribution, these values were then transformed by a chi-square quantile function, and summed) or an empirical sampling method (gene scores are transformed with chi-square quantile function and summed, then Monte Carlo estimate of the p values were obtained by sampling random sets of the same size). Results from each gene and pathway algorithm were compared within each sex as well as between sexes. Pathway information was obtained from KEGG,^20^ Reactome,^21^ and Biocarta,^22^ as made available in MSigDB.^23,24^

For genes within regions that contain SNPs previously identified as significant by GWAS, conditional analyses were run for all SNPs within those regions using SNPTEST and adjusted gene-scores were calculated. All figures were generated using R 3.3.2, ggplot2, graphite, network, Intergraph, ggnetwork, igraph, and gridExtra.^25–30^

## RESULTS

159,706 SNPs from the testing set and 163,115 SNPs from the validation set were included in gene-based analyses. Gene scores were generated for ~16,000 genes and were considered significant at p<3.3x10^−6^ (based on a Bonferroni correction for 15,000 tests). P values in the validation set were considered significant at p<0.001 (based on a Bonferroni correction for 50 tests, [25 total genes tested in each sex]).

Among males, 25 genes within five regions had scores that reached the set significance threshold (p<3.3x10^−6^) in at least 2 of 3 evaluated algorithms in all glioma or glioblastoma (See **Figure 2** and **Supplemental Table 2** for the strongest associations within each region of the six regions where genes met the set significance threshold). 19 genes within six regions had scores that reached the set significance threshold for females (p<3.3x10^−6^) in at least 2 of 3 evaluated algorithms in all glioma or glioblastoma (See **Figure 2** and **Supplemental Table 3** for the strongest associations within each of the six regions where genes met the set significance threshold). Solute carrier family 6, member 18 (*SLC6A18*), Telomerase reverse transcriptase (*TERT*), and cyclin dependent kinase inhibitor 2B (*CDKN2B*), and stathmin 3 (STMN3) reached the set significance threshold in both males and females in glioblastoma, while *SLC6A18, TERT*, and *STMN3* reached the set significance threshold in both sexes in all glioma. All shared associations validated.

**Figure 2.**
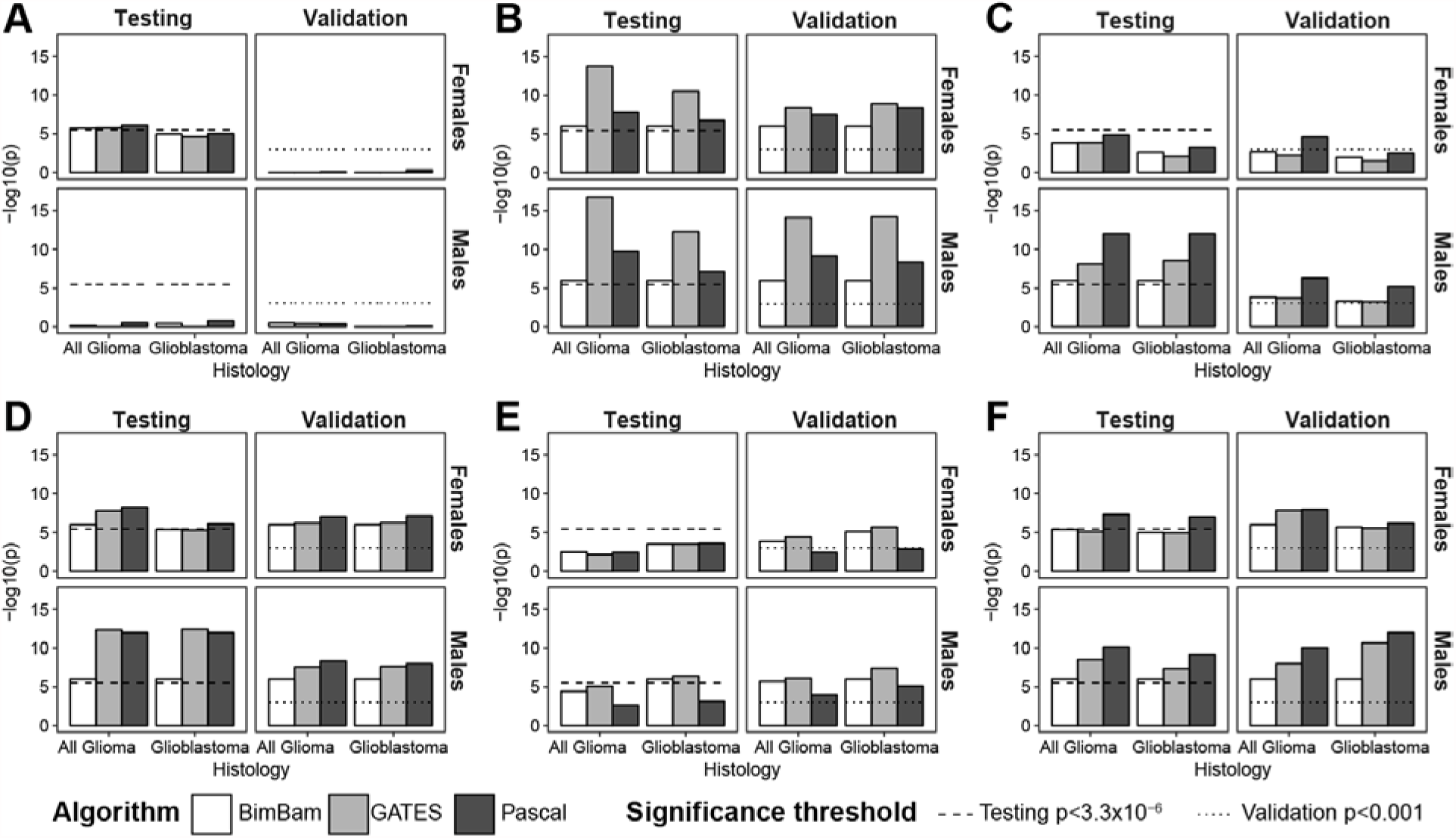
*Gene scores for prioritized genes by algorithm, histology, and sex for a) BPESC1 (3q23), B) TERT (5p15.33), C) EGFR (7p11.2), D) CDKN2B (9p21.3), E) DNAH2 (17p13.1), F) RTEL1-TNFRSF6B (20q13.33)*

Epidermal growth factor receptor (*EGFR*), dynein axonemal heavy chain 2 (*DNAH2*), and several genes surrounding regulator of telomere elongation helicase 1 (*RTEL1*) on chromosome 20 (including, RTEL1-TNFRSF6B [*RTEL1-TNFRSF6B*]) reached the significance threshold in males only (**Figure 2**). In all glioma, *CDKN2A* reached the set significance threshold in males only. All genes validated in males. Blepharophimosis, epicanthus inversus and ptosis, candidate 1 (non-protein coding) (*BPESC1*) reached the significance threshold in all glioma in females only (**Figure 2**), but this association was not able to be validated.

The association in *EGFR* was nominally significant in males after conditioning on three SNPs previously identified by GWAS within this gene (rs75061358, rs723527, and rs11979158), including one (rs11979158) that has previously been identified as having a sex-specific effect (**Supplemental Tables 4-5**). Associations in *STMN3* and *RTEL1-TNFRSF6B* were also nominally significant after conditioning in both males and females (**Figure 3, Supplemental Tables 4-5**). The association at *TERT* was nominally significant for females in glioblastoma only after conditioning on the previous identified SNP (**Figure 3, Supplemental Table 5**).

**Figure 3.**
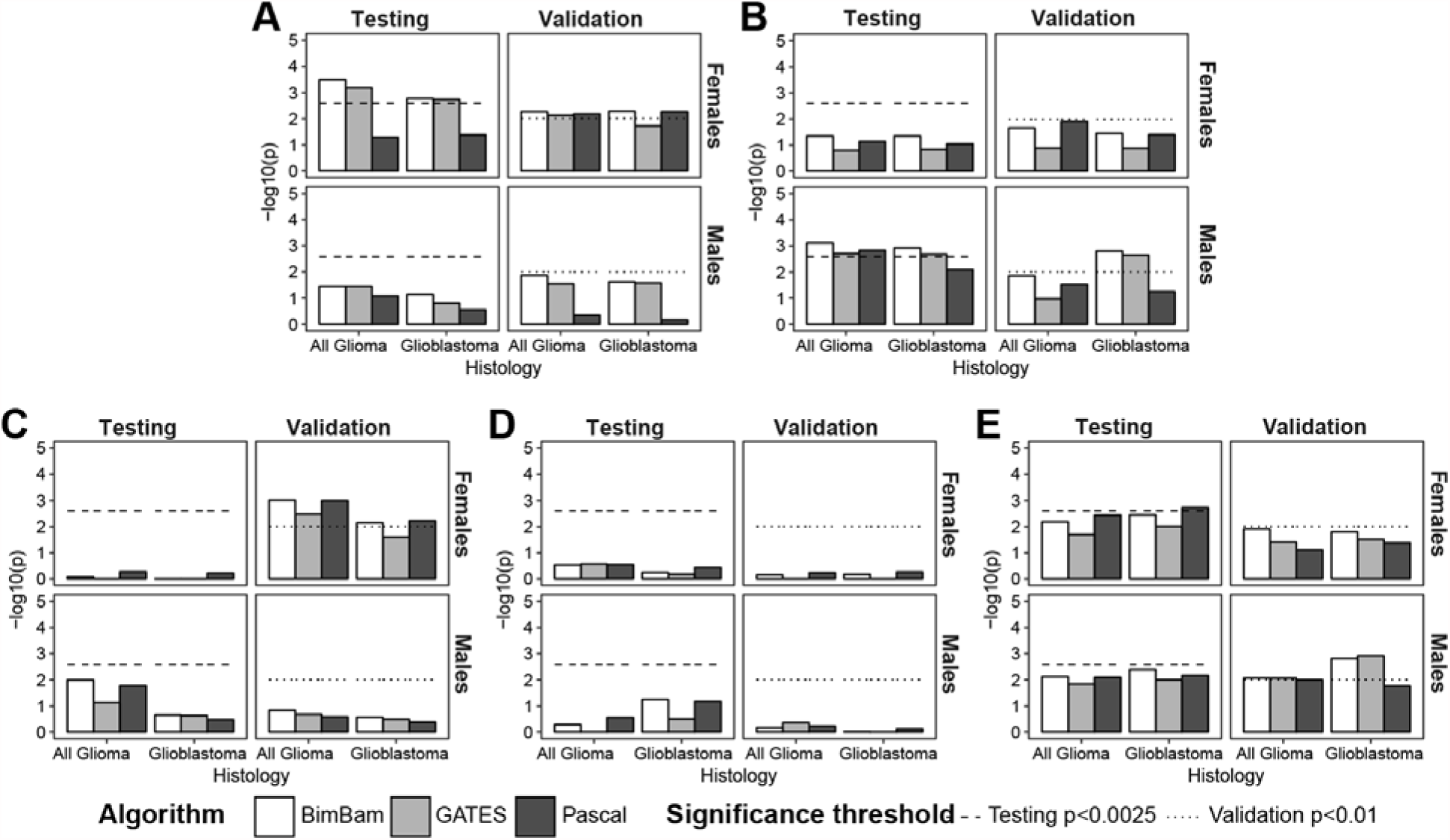
*Conditional gene scores for prioritized genes by algorithm, histology, and sex for A) TERT (5p15.33), B) EGFR (7p11.2), C) CDKN2B (9p21.3), D) DNAH2 (17p13.1), E) RTEL1-TNFRSF6B (20q13.33)*

There were 202,886 X chromosome SNPs with MAF≥0.01 and INFO score≥0.7 in the GICC dataset. Gene scores were calculated for 56 X chromosome genes with at least 5 SNPs, and associations were considered significant at p<8.3x10^−4^ (Bonferroni correction for 60 tests). There were 12 genes within 4 chromosomal regions that reached the significance threshold in at least two of three algorithms (Results from the strongest association in each region are shown in **Table 1**). Shroom Family Member 2 (*SHROOM2*) (Xp22.2), and Armadillo Repeat Containing, X-Linked 2 (*ARMCX2*) (Xq22.1) were significantly associated with both all glioma, and glioblastoma, while dystrophin (*DMD*) (Xq21.2-p21.1) was significantly associated with all glioma only, and zinc finger protein 185 with LIM domain (*ZNF185*) was significantly associated with glioblastoma only.

**Table 1.** *Gene scores for prioritized X chromosome genes by histology*

| gene (location) | histology | Pascal |  | BimBam |  |  | GATES |  |  | Algorithms<br>p<8.3x10<sup>-4</sup> |
| --- | --- | --- | --- | --- | --- | --- | --- | --- | --- | --- |
|  |  | SNPs<sup>a</sup> | P | SNPs<sup>a</sup> | Tests<sup>b</sup> | P | SNPs<sup>a</sup> | Tests<sup>b</sup> | P |  |
| SHROOM2 (Xp22.2) | All glioma | 7 | 1.20x10<sup>-4</sup> | 6 | 5.09 | 7.68x10<sup>-4</sup> | 6 | 5.09 | 0.0020 | 2/3 |
|  | Glioblastoma | 9 | 1.45x10<sup>-5</sup> | 8 | 7.06 | 5.02x10<sup>-4</sup> | 8 | 7.06 | 0.0012 | 2/3 |
| DMD (Xp21.2-p21.1) | All glioma | 88 | 3.22x10<sup>-5</sup> | 79 | 59.92 | 3.13x10<sup>-4</sup> | 79 | 59.92 | 0.0026 | 2/3 |
|  | Glioblastoma | 39 | 6.53x10<sup>-6</sup> | 37 | 31.23 | 0.0047 | 37 | 31.23 | 0.0097 | 1/3 |
| ARMCX2 (Xq22.1) | All glioma | 49 | 1.07x10<sup>-4</sup> | 44 | 33.78 | 1.89x10<sup>-4</sup> | 44 | 33.78 | 4.79x10<sup>-4</sup> | 3/3 |
|  | Glioblastoma | 63 | 5.82x10<sup>-5</sup> | 58 | 45.41 | 2.13x10<sup>-4</sup> | 58 | 45.41 | 0.0011 | 2/3 |
| ZNF185 (Xq28) | All glioma | 40 | 0.0018 | 33 | 24.26 | 0.0026 | 33 | 24.26 | 0.0061 | 0/3 |
|  | Glioblastoma | 49 | 6.19x10<sup>-5</sup> | 42 | 33.52 | 3.04x10<sup>-4</sup> | 42 | 33.52 | 9.22x10<sup>-4</sup> | 2/3 |

**Abbreviations:** *SHROOM2*: shroom family member 2; *DMD*: dystrophin; *ARMCX6*: armadillo repeat
containing, X-linked 6; *ARMCX2*: armadillo repeat containing, X-linked 2; *ZNF185*: zinc finger protein
185 with LIM domain.
a. Nominally significant (p<0.01) SNPs used in calculating gene score
b. Number of independent SNPs after filtered for linkage disequilibrium

There were 1,077 pathways in the combined KEGG, BioCarta, and Reactome sets, and associations were considered significant in the discovery set at p<5x10^−5^ (Bonferroni correction for 1,000 tests), and significant in the discovery set at p<0.0883 (Bonferroni correction for 6 tests). No pathways reached the set significance threshold, but there were several nominally significant associations. The Telomeres, Telomerase, Cellular Aging, and Immortality pathway reached nominal significance in both males and females in all glioma, and glioblastoma (**Table 2**). When the gene-scores for the genes contained within this pathway were examined, the association with this pathway was driven primarily by strong associations in *TERT*, and *TP53* (**Figure 4**). There were nominally significant associations in *POLR2A* (in both males and females) and *PRKCA* (in males only), both genes that have not been significantly associated with glioma to date. Further interrogation of the single-SNP results for these genes found no associations significant at the p<5x10^−4^ level in either sex or histology group.

**Figure 4.**
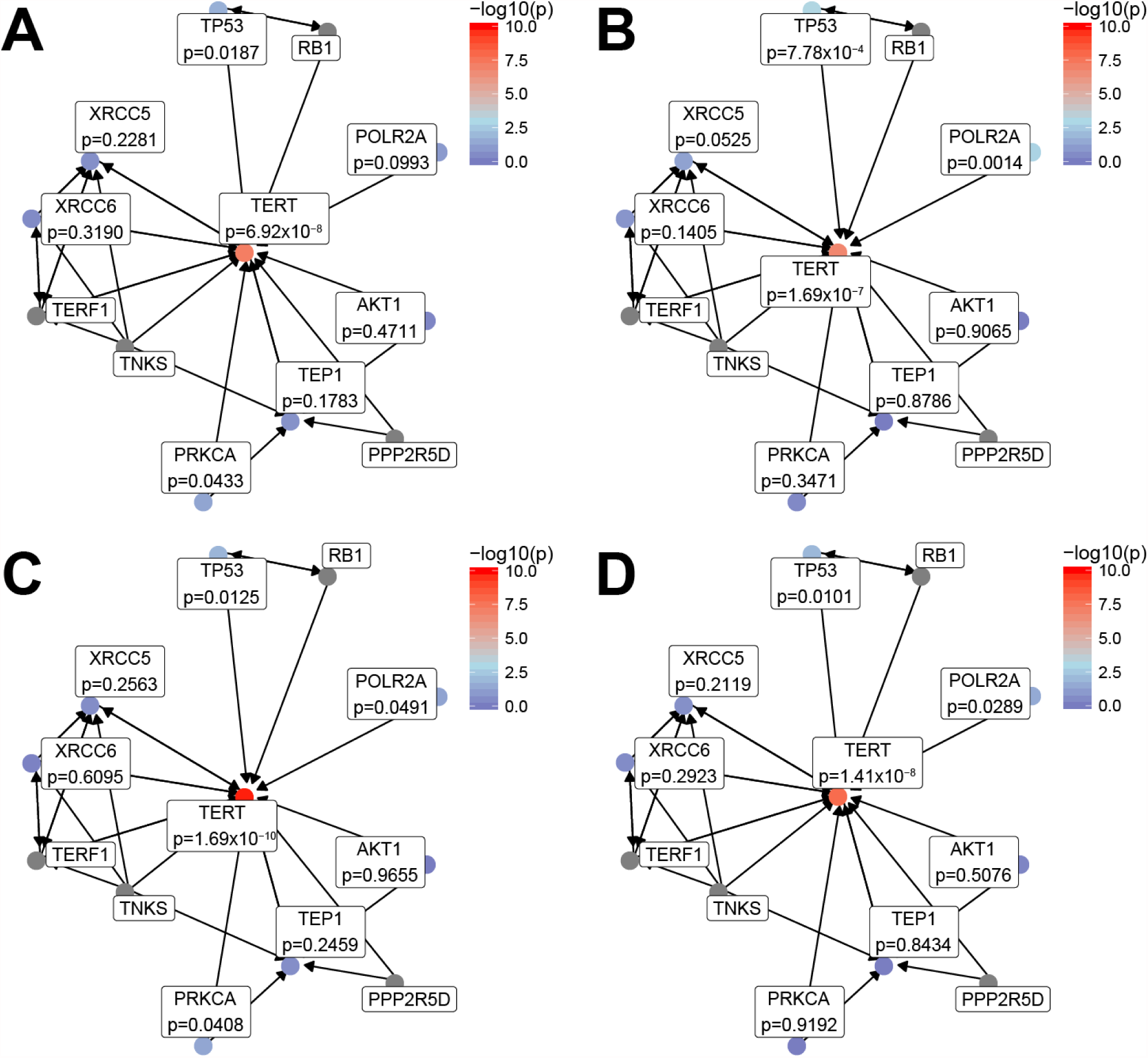
*Biocarta telomere pathway for all glioma in a) males, and b) females, and for glioblastoma in c) males and d) females*.

**Table 2.** *Significant pathways (p<0.001 in any testing group) by sex and histology*

| Pathway (Database) | Histology | Overall |  |  |  | Conditioned on previous GWAS hits |  |  |  |
| --- | --- | --- | --- | --- | --- | --- | --- | --- | --- |
|  |  | Males |  | Females |  | Males |  | Females |  |
|  |  | Discovery | Validation | Discovery | Validation | Discovery | Validation | Discovery | Validation |
| Telomeres, Telomerase, Cellular Aging, and Immortality (BioCarta) | All glioma | $5.32 \times 10^{-5}$ | $6.50 \times 10^{-4}$ | $2.61 \times 10^{-4}$ | 0.0018 | 0.3651 | 0.1046 | 0.2733 | 0.1050 |
| | Glioblastoma | $5.90 \times 10^{-5}$ | $8.60 \times 10^{-4}$ | $8.30 \times 10^{-4}$ | 0.0041 | 0.5315 | 0.4286 | 0.6885 | 0.1804 |
| Bladder cancer (KEGG) | All glioma | $9.00 \times 10^{-5}$ | 0.0013 | 0.0306 | 0.0038 | 0.5775 | 0.0514 | 0.8096 | 0.0548 |
| | Glioblastoma | $1.27 \times 10^{-4}$ | $5.50 \times 10^{-4}$ | 0.0045 | 0.0030 | 0.4097 | 0.0679 | 0.7187 | 0.0394 |
| Glioma (KEGG) | All glioma | $5.80 \times 10^{-4}$ | 0.0057 | 0.0361 | $5.00 \times 10^{-4}$ | 0.5595 | 0.1241 | 0.5701 | 0.0045 |
|  | Glioblastoma | 0.0011 | 0.0061 | 0.0048 | 0.0018 | 0.4818 | 0.2616 | 0.3783 | 0.0132 |
| Melanoma (KEGG) | All glioma | $7.60 \times 10^{-4}$ | 0.0032 | 0.0219 | $1.70 \times 10^{-4}$ | 0.7131 | 0.0650 | 0.5803 | 0.0016 |
| | Glioblastoma | $8.70 \times 10^{-4}$ | 0.0020 | 0.0013 | $7.60 \times 10^{-4}$ | 0.5468 | 0.0934 | 0.2391 | 0.0054 |
| Non-small cell lung cancer (KEGG) | All glioma | $7.30 \times 10^{-4}$ | 0.0171 | 0.0290 | $7.40 \times 10^{-4}$ | 0.7866 | 0.4067 | 0.6097 | 0.0067 |
|  | Glioblastoma | 0.0013 | 0.0455 | 0.0011 | 0.0019 | 0.6823 | 0.8232 | 0.2093 | 0.0174 |
| Pancreatic cancer (KEGG) | All glioma | $2.65 \times 10^{-4}$ | 0.0132 | 0.0021 | 0.0016 | 0.3903 | 0.2530 | 0.0957 | 0.0133 |
| | Glioblastoma | 0.0021 | 0.1124 | $2.01 \times 10^{-4}$ | $9.10 \times 10^{-4}$ | 0.6711 | 0.9026 | 0.0413 | 0.0060 |

Nominally significant associations were identified in 5 cancer-specific KEGG pathways: bladder cancer, glioma (**Supplemental Figure 2**), melanoma (**Supplemental Figure 3**), non-small cell lung cancer, and pancreatic cancer (**Table 2**). There is significant overlap between these gene-sets (**Supplemental Figure 4**), and when the gene scores used to build each pathway were examined all the associations appear to be driven largely by strong associations in *EGFR,* and *CDKN2A* which are members of all KEGG cancer pathways found to be nominally associated with glioma in this analysis. Pathway analyses were run using single-SNP results including conditional analyses for all SNPs within a 2mb window around the previously identified SNPs in the *TERT, EGFR, CDKN2B, TP53* and *RTEL1* loci. All pathway associations no longer reached the significance threshold when analysis included conditioned results (**Table 2**).

DISCUSSION

Multi-marker tests, such as gene- or pathway-based tests, allow investigators to leverage previously existing summary statistics and increase power when strength of single-SNP associations may be low. This analysis aimed to explore additional potential sources of genetic risk that may contribute to sex differences in genetic risk for glioma. All autosomal genes identified by and validated within this analysis were proximate to previously identified GWAS hits. After conditioning on these previously identified SNPs, regions including *TERT, EGFR* and *RTEL1* remained nominally significant, while associations at the other identified genes were no longer significant. While GWAS has identified one locus near *TERT*, two independent loci near *EGFR*, and one loci near *RTEL1* that are highly significantly associated with glioma risk, the results of this conditional analysis suggest that there are remaining sources of genetic risk for glioma within these regions.

Four regions on the X chromosome (Xp22.2, Xp21.2-p21.1, Xq22.1, and Xq28) contained genes that reached the significance threshold in at least two of three algorithms (**Table 1**). None of these genes have been previously associated with glioma. SNPs surrounding *SHROOM2* (Xp22.2) were previously associated with prostate and colon cancer.^31–33^ There are no known associations with inherited variants in the other four regions and increased risk for cancer, though all have been shown to be dysregulated in some cancer cells. *DMD* encodes for dystrophin, which is an essential component of muscle tissue. Inherited or de novo mutations in *DMD* are well known to cause a spectrum of muscle diseases called dystrophinopathies (including Duchenne muscular dystrophy, and Becker muscular dystrophy).^34^ Deletions in this gene have been found in mesenchymal and stromal tumors, and downregulation of this gene has been associated with progression and metastasis in these tumors.^35,36^ *ARMCX2* (Xq22.1) is a member of the armadillo family of proteins, several of which have been implicated in tumorigenesis.^37^ *ARMCX2* has been shown to be differentially expressed in cancer cell lines as compared to normal cell lines, though expression in glioma cell lines does not differ from normal.^38^ The protein encoded by this gene has been shown to be decreased in lung cancer and expression of *ZNF185* is negatively correlated with progression in prostate cancer where it is silenced by methylation.^39,40^Without a validation set, it is not possible to know if these are true associations or the result of type 1 error. Further exploration of these genes is necessary to determine their true relationship with glioma risk.

The *Telomeres, Telomerase, Cellular Aging, and Immortality* pathway reached nominal significance in both males and females in all glioma, and glioblastoma (**Table 2**). This pathway contains *EGFR, TERT*, and *TP53*, all of which contain variants that have been previously associated with increased odds of developing glioma. Variants associated with telomere maintenance have been associated with glioma, as swell as many other complex diseases.^41–;43^ An analysis comparing a weighted genetic score based on 8 SNPs associated with leukocyte telomere length (LTL) (*ACYP2, TERC, NAF1, TERT, OBFC1, CTC1, ZNF208*, and *RTEL1*) found that telomere length was ~5% longer in glioma cases versus controls.^44^ The significance of the telomere maintenance pathway may explain the remaining significant association in the regions surrounding *TERT, EGFR* and *RTEL1*, as any variants affecting telomere length could contribute to glioma risk. In addition to the strong associations in genes associated with SNPs previously identified by glioma GWAS, there were nominally significant associations in *POLR2A* (in both males and females) and *PRKCA* (in males only).

The numerous KEGG cancer pathways found to be significant in this analysis are likely due to the strength of association in genes (*CDKN2A, EGFR*) that are members of many pathways. While these associations are driven by these specific genes, they may also be evidence of shared genetic pathways in sources of genetic risk, or process of carcinogenesis between these cancers and glioma. Both the KEGG glioma and melanoma pathways were significantly associated with all glioma in males, both of which appear to be strongly driven by associations in *CDKN2A* (**Supplemental Figures 3-4**). Previous analyses suggested an association between genetic risk for glioma and melanoma, both in terms of syndromic cancer (most notably Melanoma-neural system tumor syndrome, caused by inherited variants in *CDKN2A*^2^), familial glioma and sporadic disease. An analysis of the NCI’s SEER system found that persons with a previous diagnosis of melanoma have incidence of glioma that is 1.42x that of the general population.^45^ Family based studies have found that relatives of glioma patients have higher than expected incidence of melanoma, approximately 2-4 times that of the general population.^46,47^Genome-wide association studies for melanoma to date have identified at least 21 genetic risk loci.^48,49^including SNPs in the regions surrounding *CDNK2A* and *TERT* that have been previously associated with glioma.^4^ These SNPs do not account for a large proportion of risk in either cancer type, but there is some evidence that telomere length and pathways of telomere maintenance may contribute to risk in both diseases ^50^.

When pathway analysis were re-run using conditioned single-SNP results, pathway associations no longer reached the significance threshold when analysis included conditioned results. Gene-specific p values for conditional analyses of *TERT, EGFR*, and *RTEL1* were lowest in analyses performed in Pascal, which are the gene-scores used in calculated the pathway-specific results. Pascal uses a SOCS approach, which may be conservative than others if there are many SNPs with null association and few SNPs with significant associations. If this is the case in this

While multi-marker tests (including gene, and pathway tests) have the ability to increase power to detect associations as compared to single-SNP tests, different methods will perform differently and may be better suited to particular types of genetic architecture. Results for methods that use LD information, including all algorithms evaluated in this analysis, may also be significantly altered by the reference populations to estimate LD. All of the included methods attempt to adjust for potential score inflation due to LD, using the 1,000 EUR super population as a reference set. FAST does this by pruning the data of SNPs that are in complete linkage (r^2^=1), while Pascal does this by generating ‘fusion’ gene scores for genes that are in linkage with each other. These ‘fusion’ genes are then utilized in pathway analyses to avoid inflation due to the physical proximity of genes, and decreased p value inflation.^9^ Due to variations in adjustment for linkage disequilibrium used in the two programs, the number of included SNPs by each gene varied slightly. Both methods require that each variant in the summary statistics be present in the LD reference file, and as a result these methods are not able to incorporate variants that do not have a standard ID. FAST additionally limits the dataset by requiring that all markers be bi-allelic SNPs, and does not accept indels.

Both Pascal (which is an implementation of the VEGAS scoring system) and the GATES method within FAST do not rely on permutations for estimating p values. The VEGAS algorithm as implemented within FAST,^19^ and VEGAS2^51^ both rely on Monte Carlo simulations to estimate P values. Permutation-based tests are significantly more computationally intensive, especially when gene scores are being calculated across the entire genome. BimBam uses permutations to calculate exact p values, as a result is more computationally intensive. The number of permutations used to calculate determines the boundaries for an exact p value (ranging from 1 to 1/n, where n is the number of permutations), which may result in increasing permutations for increased p value specificity. Pascal allows for both a sum of chi square, as used in this analysis, or a maximum chi square calculation of the test statistics. All of these methods require consideration of the assumptions being made about the genetic architecture of the disease and population of interest.

There is a well-known bias in GWAS towards large genes,^52^ and this bias may influence the results of this analysis. Large genes may be enriched for tag SNPs selected on arrays, and will be further enriched through imputation. All of the algorithms used for this analysis can be effected by gene size. Large genes with many SNPs of minimal significance and few SNPs of large effect may ‘dilute’ the gene score in methods based on summed scores, such as Pascal and VEGAS. All algorithms prune SNPs in attempt to obtain a set of independent SNPs, but this may still bias results towards large genes if the gene contains multiple haplotype blocks. This analysis used a relatively large window surrounding the defined genes (+/- 50kb) which may further bias analyses towards large genes.

This represents the first genome-wide sex-specific gene- or pathway-based analysis for germline risk variants in glioma. Gene-based tests are an efficient way to increase power to detect associations of low effect size, where multiple variants within a region may contribute to increased risk. This analysis provides additional support for a mechanistic association between telomere function, and glioma risk. There are several limitations to this analysis. All glioma cases from the included four GWAS datasets were recruited at time of first diagnosis, and the assigned diagnoses represent the primary tumor type. There may also be variation in the histologies contained within each set by sex. The proportion of each dataset that is composed of glioblastoma as compared to lower grade gliomas varies by both study and sex (**Supplemental Table 1**). Less than 50% of female glioma cases in the testing set are glioblastoma, whereas over 50% of female cases are glioblastoma in the validation sets. Glioma is a heterogonous disease, and due to all of these factors, it is likely that heterogeneity exists between the utilized datasets.

## CONCLUSIONS

Multi-marker tests, such as gene- or pathway-based tests, allow investigators to leverage previously existing summary statistics and increase power when strength of single-SNP associations may be low. This analysis aimed to explore additional potential sources of genetic risk that may contribute to sex differences in genetic risk for glioma. There was a nominally significant association between germline variants in *RTEL1* in both males and females after conditioning on previously identified SNPs. There was also a significant association between germline variants in the telomere maintenance pathway in both males and females, which builds on previous evidence of the relationship between inherited variants related to increased telomere length and increased risk for glioma. There was also a male specific association in *EGFR*, and a female-specific association in *TERT* which remained nominally significant after conditioning on previous GWAS hits. The results of this analysis confirm previously known information about inherited glioma risk, and provide potential mechanistic explanations for how these variants may affect the process of gliomagenesis.

## Acknowledgments

Other significant contributors for the UCSF Adult Glioma Study include: M Berger, P Bracci, S Chang, J Clarke, A Molinaro, A Perry, M Pezmecki, M Prados, I Smirnov, T Tihan, K Walsh, J Wiemels, S Zheng. Glioma scan group comprised: Laura E. Beane Freeman, Stella Koutros, Demetrius Albanes, Kala Visvanathan, Victoria L. Stevens, Roger Henriksson, Dominique S. Michaud, Maria Feychting, Anders Ahlbom, Graham G. Giles Roger Milne, Roberta McKean-Cowdin, Loic Le Marchand, Meir Stampfer, Avima M. Ruder, Tania Carreon, Goran Hallmans, Anne Zeleniuch-Jacquotte, J. Michael Gaziano, Howard D. Sesso, Mark P. Purdue, Emily White, Ulrike Peters, Howard D. Sesso, Julie Buring. We are grateful to all the patients and individuals for their participation and we would also like to thank the clinicians and other hospital staff, cancer registries and study staff in respective centers who contributed to the blood sample and data collection.

